# Bile-mediated ToxS homodimerization provides a model for ToxRS periplasmic interactions

**DOI:** 10.64898/2026.01.06.697910

**Authors:** Minje Kim, Deepak Balasubramanian, F. Jon Kull, Salvador Almagro-Moreno, Charles R. Midgett

## Abstract

The ToxRS system belongs to a family of co-component transmembrane transcription regulators that act as sensors of environmental cues and regulate virulence gene expression in several bacterial pathogens. These systems are thought to operate by sensing environmental stimuli and transmitting signals through periplasmic domains to activate DNA-binding transcription factors. In the enteric pathogens *Vibrio parahaemolyticus* and *Vibrio cholerae*, the ToxRS system regulates virulence factors responsible for severe gastrointestinal symptoms in humans. ToxR is a DNA-binding regulator associated in the periplasm with ToxS, a protein of poorly understood function. ToxS modulates the activity of its binding partner ToxR in the presence of bile salts, antimicrobial cholesterol metabolites secreted into the gut. To date, the molecular mechanism underlying this regulation remains unclear. We present crystal structures of the *V. parahaemolyticus* ToxS periplasmic domain (ToxSp) with and without the bile salt glycocholate. ToxSp forms an 8-stranded broken β-barrel with a central α-helix and is structurally homologous to a group of chaperone proteins. ToxSp has a highly conserved hydrophobic core that stabilizes the β-barrel fold, while the binding pocket tolerates substantial variation, consistent with binding hydrophobic ligands. Strikingly, we discovered that Vp-ToxSp binds three molecules of glycocholate and the presence of this bile salt leads to the formation a strand-swapped ToxS homodimer. Finally, modeling two ToxR periplasmic domains in complex with the glycocholate-bound ToxSp homodimer provides a structure-based model for bile salt-mediated heterotetramerization of the ToxRS system. Overall, our study addresses a major longstanding question in the field of Vibrio virulence regulation providing a scenario that could apply to other pathogens that utilize these membrane-bound family transcriptional regulators.

**AUTHOR SUMMARY:** The ability to sense the environment is essential for bacterial pathogens to successfully colonize the human host. Many pathogenic bacteria utilize members of the family of co-component transmembrane transcription regulators (coTTRs) to regulate pathogenesis in response to host signals. The ToxRS system is a coTTR regulatory pair that is essential for host colonization and virulence regulation of the enteric pathogens *Vibrio cholerae* and *Vibrio parahaemolyticus*. This system encompasses a transcription factor, ToxR, and a binding partner of enigmatic function, ToxS. Here, we solved the structure of the ToxS periplasmic domain *<u>V. parahaemolyticus</u>* and found that it belongs to a larger family of chaperone-like proteins present in many organisms. Strikingly, we discovered that the presence of the bile salt glycocholate leads to the dimerization of the periplasmic domain of ToxS. We propose that bile salt-induced ToxS dimerization brings two ToxR cognates together forming a heterotetramer activating the regulatory system. Our results and model address a major longstanding question in the fields of Vibrio virulence regulation and could apply to other pathogens that utilize these membrane bound family transcriptional regulators.

## INTRODUCTION

Many pathogenic bacteria utilize the family of co-component transmembrane transcription regulators (coTTRs) to regulate pathogenesis in response to host signals [1]. These include VtrAC and ToxRS in *Vibrio* spp., PsaEF from *Yersinia* spp., GrvA-FidL from enterohemorrhagic *Escherichia coli*, and YqeIJ from avian pathogenic *E. coli* [2–8]. CoTTRs consist of a transmembrane transcription factor with an N-terminal winged helix-turn-helix DNA-binding domain, a transmembrane helix and a C-terminal periplasmic domain, together with a periplasmic binding partner that has a short N-terminal tail, a transmembrane helix, and a C-terminal periplasmic domain [1,9]. While the periplasmic domains of these proteins have little sequence homology between family members, many are clearly structural homologs [7,10,11]. These two-protein systems form obligate heterodimers where they modulate gene expression in response to host signals for survival and colonization [1,12]. This is thought to occur by the binding partner sensing a stimulus, such as a small molecule, and then transducing the signal to the transcription factor through their respective periplasmic domains [7,12,13]. To date, the mechanistic link between signal sensing by the periplasmic binding partner and DNA binding by the cognate transcription factor remains unknown.

The ToxRS coTTR system acts as an environmental sensor and is conserved across the Vibrionaceae family, a diverse group of aquatic bacteria [14–16]. Importantly, in several pathogenic *Vibrio* species, ToxRS also acts as a virulence regulator and confers resistance against antimicrobial insults found in the GI tract such as bile salts, cholesterol metabolites produced in the liver and secreted into the intestine [3,15,17,18]. In *Vibrio cholerae,* the etiological agent of cholera, ToxRS helps mediate the stress response to bile salts by modulating the transcription of two outer membrane porins, OmpU and OmpT [19,20]. Furthermore, together with another coTTR pair, TcpPH, ToxRS activates the virulence cascade via the transcription of the master virulence regulator *toxT* [21–23]. ToxT directly promotes the production of two critical virulence factors: the cholera toxin and toxin co-regulated pilus, which are essential for disease symptoms to occur [24–27]. Similarly, in *Vibrio parahaemolyticus*, a source of gastrointestinal illness, ToxRS regulates bile responses and essential virulence genes [3,18]. Together with another coTTR system, VtrAC and ToxRS promote the expression of *vtrB*, which activates transcription of genes encoding the type III secretion system [3,28,29].

Numerous studies have demonstrated the role of ToxR in these processes. However, despite its relevance and ubiquitousness, the specific role of ToxS has remained obscure for decades. Studies using mouse models of *V. cholerae* and *V. parahaemolyticus* infection show that ToxS is essential for intestinal colonization, with Δ*toxS* strains exhibiting severe colonization defects [3,30]. Our laboratories and others have shown that ToxS stabilizes ToxR against premature intramembrane proteolysis by leading to a conformation that induces the formation of an intra-chain disulfide bond [31–33]. Furthermore, ToxS is required for full ToxR activity in response to several stimuli including bile salts [13,34,35]. In minimal media, ToxR requires ToxS to respond to bile salts and the amino acid mixture NRES [36,37]. Overall, these data suggest that ToxRS exhibits on and off states, with ToxS being central for this switch-like process [36,37]. To date, the molecular mechanisms that mediate ToxRS interactions and the specific role of ToxS in these processes remain unknown. Here, we solved the structure of the apo and glycocholate-bound ToxS periplasmic domain (ToxSp) from *V. parahaemolyticus* (Vp-ToxSp). ToxSp is monomeric in the apo state and dimeric in the bile salt-bound state, suggesting ToxS directly senses environmental stress signals and dimerizes upon ligand binding. Additionally, a structural analysis of ToxSp suggests it is related to a group of chaperone proteins. Our results suggest that coTTRs oligomerization is directed by interactions of the periplasmic binding partner with environmental stimuli, which leads to heterotetramer formation with the cognate transcription factor.

## RESULTS

### ToxS periplasmic domain adopts a broken β-barrel fold with a putative ligand-binding pocket

The *V. parahaemolyticus* ToxS periplasmic domain residues Ser25-Asn171 (Vp-ToxSp) was cloned into an *E. coli* expression system and purified using chitin-based affinity purification. Diffraction quality apo ToxSp crystals were obtained using a seeded crystallization method and resolved to 1.9 Å resolution (**Table S1**). The resulting structure reveals the Vp-ToxSp domain adopts a monomeric eight-stranded broken β-barrel, with two copies in the asymmetric unit. The domain begins with the N-terminal helix (α1), positioned at the base of the β-barrel, proximal to the inner membrane, followed by five β-strands (β1-β5), a second α-helix (α2), and three final β-strands (β6-β8), with several disordered loops connecting the β-strands. Barrel closure is enabled by hydrogen-bonding interactions between β1 of the N-terminus and the C-terminal β-strands (β6-β8). (**Figure 1A, 1B**).

**Figure 1.**
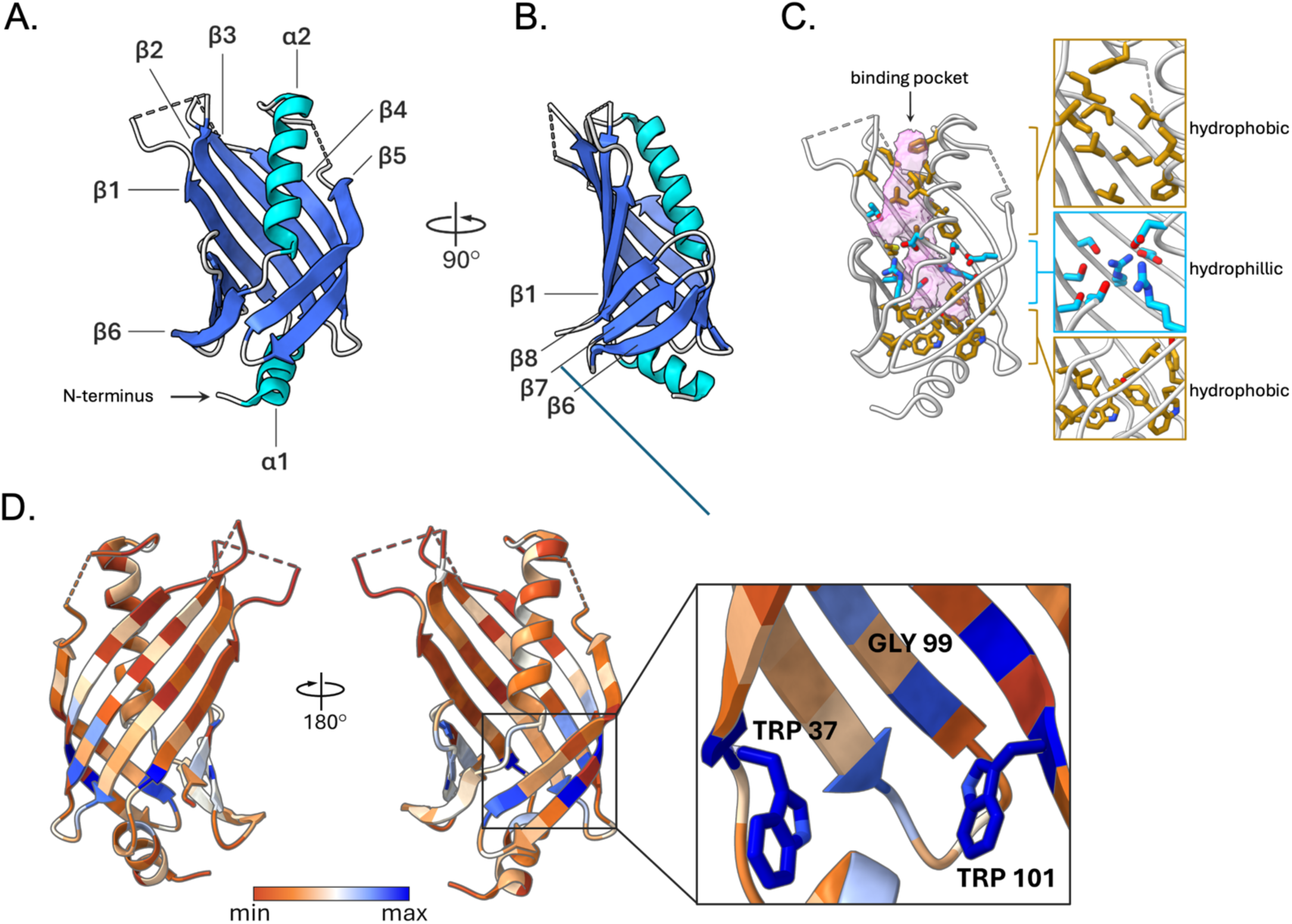
Structure and conservation of the *Vibrio parahaemolyticus* ToxS periplasmic domain. (A, B) Apo ToxSp from *V. parahaemolyticus* contains an 8-stranded broken beta-barrel fold and the central alpha-helix (β-strands; dark-blue, α-helices; light blue, coils; gray). (C) Left panel: apo ToxSp has a binding pocket (left panel, pink) enclosed by the β-barrel and the central helix (grey). Right panel: the binding pocket constitutes a central region of hydrophilic residues (blue) flanked by regions with hydrophobic (gold) residues. (D) Apo-Vp ToxSp structure colored based on conservation data obtained from sequence alignment across 280 Vibrionaceae family members reveals that only residues Trp37, Gly99, and Trp101 are 100% conserved.

The central core consists of alternating hydrophobic and hydrophilic regions and the membrane-distal face contains a solvent-accessible ∼745 Å^3^ cavity that likely functions as a small molecule binding region extending toward the membrane (**Figure 1C right**) [38]. It is characterized by hydrophobic residues forming the upper rim, a ring of hydrophilic residues in the middle, and a deeper hydrophilic cavity sealed at its base by α1 (**Figure 1C left**). Interestingly, the pocket architecture resembles those observed in previously characterized bile-binding proteins, suggesting a similar ligand-recognition role for Vp-ToxSp [7,11,39].

### Core conservation and pocket variability in ToxS

To examine the amino acid conservation across the Vibrionaceae family, 280 ToxR sequences were aligned using Unipro UGENE, and the corresponding ToxS sequences were arranged in the same order [40]. Both sets of sequences were analyzed using Clustal Omega (EMBL) to generate percent identity matrices (**Figure S1, S2**) [41]. Comparison of these matrices revealed two key trends: first the ToxS periplasmic domain (ToxSp) is more conserved than the ToxR periplasmic domain and second the clustering patterns observed for ToxSp mirror those of ToxRp, suggesting coordinated evolutionary pressures on both components of the ToxRS pair. To examine residue-level conservation, the ToxSp alignment was analyzed in ChimeraX using AL2CO [42]. The most strongly conserved residues localize to the hydrophobic core, with three residues, Trp37, Gly99, and Trp101, showing 100% conservation across all analyzed sequences (**Figure 1D and inset**). The two tryptophan residues are packed deep within the hydrophobic core, while Gly99 appears to contribute to a kink in β4, influencing the positioning of Trp101, highlighting the importance of this region in the overall structure. In contrast, residues surrounding the ligand-binding pocket exhibit low sequence identity across Vibrionaceae (**Figure 1D**). Despite this variability, many positions within the pocket retain a strong preference for hydrophobic side chains, with several sites remaining >90% hydrophobic and others maintaining hydrophobicity in ∼70% of sequences (**Table S2**). Together, these observations indicate that ToxS is built upon a highly conserved hydrophobic core that stabilizes the β-barrel fold, while the pocket tolerates substantial variation, consistent with binding hydrophobic ligands through largely non-specific interactions. Finally, low conservation at the ToxS-ToxR binding interface supports the idea that these two proteins have coevolved at their binding interfaces.

### The ToxS periplasmic domain fold is related to a large group of chaperones

Having defined the overall structure of Vp-ToxSp, we next sought to determine whether this architecture resembles any previously characterized protein families. Because structural homology can provide functional insights in the absence of primary sequence conservation, we used DALI to identify proteins with related three-dimensional folds (**Table S3**) [43]. This search revealed VtrC, its closest known homolog in *V. parahaemolyticus*, as the top hit, validating the approach. Additional significant matches containing this fold were identified, including periplasmic chaperones HslJ (PDB ID: 2KTS), first identified in *Escherichia coli* and MxiM from *Shigella flexneri* [44–46]. Structural similarity was also observed with an AlphaFold-predicted structure of META1, part of the META family in *Leishmania* spp. (**Figure 2A**), indicating that ToxS-like proteins are distributed in a wide range of taxa [47]. Despite their evolutionary diversity, these proteins share topological features, including the C-terminal β-strands (β6-β8) (green) and the central α-helix (orange) (**Figure 2B**).

**Figure 2.**
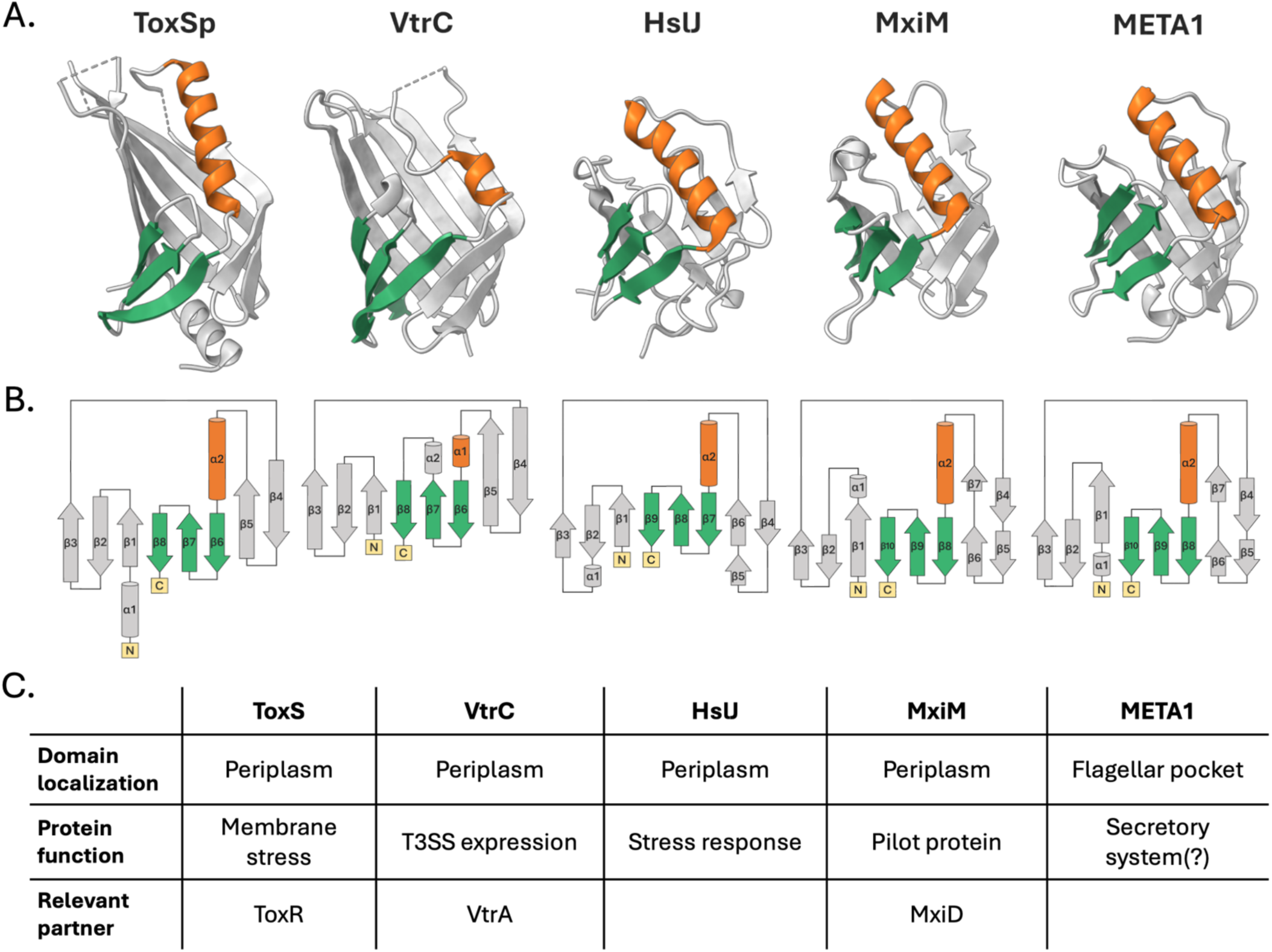
ToxSp is a stress-responsive periplasmic chaperone. (A) Structural comparison of ToxSp with VtrC (PDB ID: 5KEV), HslJ (PDB ID: 2KTS), MxiM (PDB ID: 1Y9L), and META1 (AlphaFold: A0A088S6F1) highlights a shared broken β-barrel fold common among chaperone proteins. (B) The structural and topological similarities are highlighted by the C-terminal β-strands (β6-β8) (green) and the central α-helix (orange). (C) Under each topology are relevant highlights of the protein localization, function, and relevant binding partner.

The structural and topological relatedness between these proteins is mirrored by parallels in function, as many of the identified hits serve as periplasmic chaperones and/or respond to stress. Consistent with this, ToxS stabilizes the ToxR periplasmic domain and promotes disulfide bond formation [33,48]. Likewise, VtrC in *V. parahaemolyticus* stabilizes VtrA to turn on the expression of *vtrB* that activates the type III secretion system 2 [7]. HslJ functions as a stress-induced periplasmic chaperone in multiple bacterial species [46,49–52] and MxiM acts as a pilot protein required for type III secretion system-mediated host cell invasion in *Shigella* spp. [53]. Although the role of META1 is less clear, its overexpression increases virulence in mice infected with *Leishmania amazonensis*, and the related META2 protein is upregulated in response to heat shock and oxidative stress [47,52,54,55]. Despite these structural and functional similarities (**Figure 2A**), ToxS, HslJ, MxiM, and META1 display very limited sequence homology, suggesting strong evolutionary pressure to conserve fold and function rather than the primary sequence [47].

### Vp-ToxSp bound to bile salts forms a strand-swapped dimer

The 1.9 Å crystal structure of Vp-ToxSp bound to glycocholate reveals a pronounced ligand-induced conformational change relative to the apo structure (**Figure 3A; Table S1**). Although each monomer retains the characteristic 8-stranded broken β-barrel fold with two α-helices, the bile salt-bound form assembles into a strand-swapped dimer in which β8 strands of the two monomers exchange positions (**Figure 3B**), with two dimers in the asymmetric unit. Notably, the swapped β8 strands align in the same register as in the apo state, preserving the hydrogen-bonding pattern of the β-barrel. In contrast to the apo structure, all loops at the top of the barrel are fully resolved, with the loop between β1 and β2 folding over the top of the barrel, capping the ligand-binding pocket. The orientation of the two N-termini within the dimer is compatible with the insertion of adjacent transmembrane helices into the membrane (see below).

**Figure 3.**
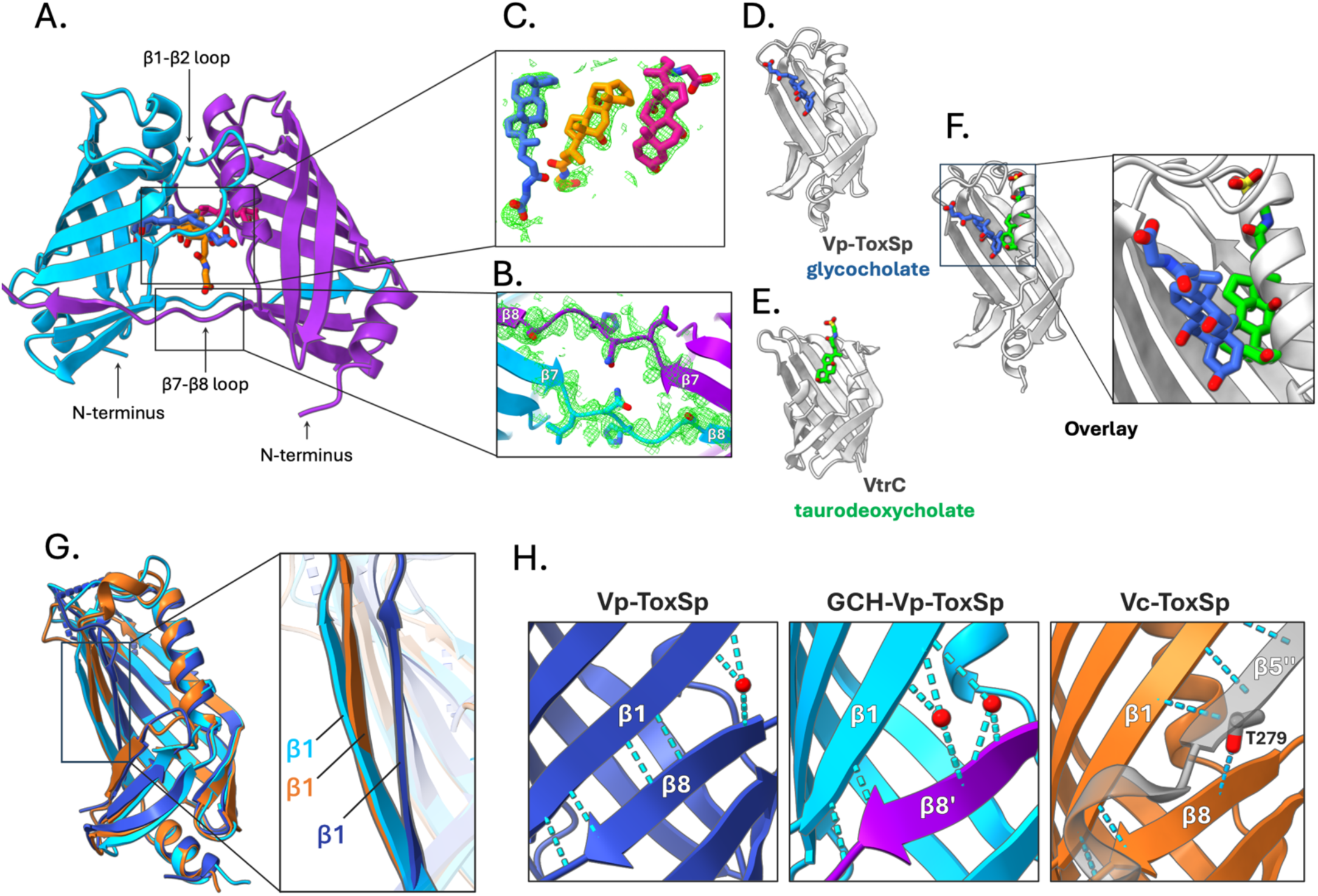
Bile salt binding drives strand swapping and homodimerization of ToxS. (A) The GCH bound ToxSp dimer crystal structure shows two β8 strand swapped ToxSp monomers (blue and purple β-barrels) enclosing three glycocholate molecules (blue, orange, and pink). (B) Electron density map of the crystal structure surrounding the strand-swapped β8 loops (Ser160-Thr166) displayed as green mesh, contoured at 1σ. (C) Electron density map of the three glycocholate molecules bound to the ToxSp dimer displayed as green mesh, contoured at 1σ. Comparison of (D) Vp-ToxSp-GCH with the protein in gray and the ligand in blue, and (E) VtrC-taurodeoxycholate (PDB ID: 5KEW) with the protein in blue and the ligand in green, complexes demonstrates the different binding orientations of the two ligands. (F) Overlay of the crystal structures of GCH-bound Vp-ToxSp monomer, and taurodeoxycholate bound to VtrC (protein not shown for clarity) highlights the differences in the ligand orientation. (G) Overlay of the apo Vp-ToxSp, (blue), and a GCH-Vp-ToxSp monomer (cyan), and the Vc-ToxSp from the heterodimeric ToxRp-ToxSp complex (orange; PDB: 8ALO). The comparison (inset) shows that apo Vp-ToxSp (blue) adopts a more closed conformation, whereas both glycocholate-bound Vp-ToxSp (cyan) and Vc-ToxSp (orange) from the heterodimer with ToxR display a more open conformation. (H) In apo Vp-ToxSp (blue, left), a single water molecule (red) bridges β1 and β8 strands to stabilize the closed conformation. In GCH-bound Vp-ToxSp (cyan, middle), two water molecules (red) bind the more open interface, whereas in Vc-ToxSp (orange, right), Thr279 from Vc-ToxRp (grey) plays the role of the water molecules, stabilizing the closed conformation.

### Three glycocholate molecules stabilize the ToxSp dimer interface

The strand-swapped ToxSp dimer structure contains three glycocholate molecules bound between the two monomers toward the top of the β-barrel. In each monomer, glycocholate binds the hydrophobic pocket formed by residues from β1, β2, β3, and α2 (**Figure S3A**). The third glycocholate occupies a central site created by the juxtaposition of the C-terminal ends of two α2 helices from the two monomers, effectively bridging the two monomer-bound ligands. Across these sites, the glycocholates adopt a common orientation: their hydrophobic rings embed into the upper hydrophobic region of the binding pocket, while the polar hydroxyl and glycine groups remain solvent-exposed (**Figure 3C**). Although the interaction network is broadly conserved, the monomers contribute asymmetrically to ligand contacts at the interface. Two additional glycocholate molecules are present at the crystal-packing interfaces but display weaker electron density and are unlikely to be physiologically relevant (**Figure S3B**). The presence of an extended binding pocket in the ToxSp dimer that is absent in the apo structure supports a model in which binding of three glycocholate molecules promotes ToxSp dimerization. β8 strand swapping further stabilizes this conformation. These observations indicate that ToxS functions as a ligand-responsive sensor that undergoes a monomer-to-dimer transition upon bile binding.

### ToxS and VtrC bind bile salts through distinct mechanisms

Comparison of bile salt-bound structures of ToxSp and VtrC demonstrates that, despite their shared broken β-barrel architecture, these proteins bind bile salts in distinct ways. In VtrC, taurodeoxycholate and chenodeoxycholate occupy a pocket formed directly within the β-barrel [7,39]. In ToxSp, however, helix α2 occupies the position taken up by bile salts in VtrC, and the β1-β2 loop folds over the top of the barrel, an architectural feature that is absent in VtrC (**Figure 3D, 3E**). As a result, the orientation of glycocholate differs substantially: the side chain of glycocholate projects outward, rather than upward, as observed in VtrC (**Figure 3F**). This outward orientation exposes the hydrophilic tail of the glycine moiety to solvent, potentially allowing ToxSp to accommodate bile salts with diverse side chains. These differences suggest that although ToxS and VtrC share an overall broken β-barrel fold, the coTTR binding partners have evolved distinct ligand-binding strategies while preserving the conserved broken β-barrel scaffold.

### ToxSp bound to bile adopts a conformation resembling the ToxR-bound state

To determine whether bile salt binding induces conformational changes relevant to ToxR engagement, we compared the apo and glycocholate-bound *V. parahaemolyticus* ToxSp structures with the *V. cholerae* ToxSp (Vc-ToxSp) from the heterodimeric ToxRS complex. This analysis reveals that bile salt binding drives Vp-ToxSp into a conformation closely resembling the ToxR-bound state (**Figure 3G**) [11]. In the apo structure of Vp-ToxSp, the β1 strand of the barrel is bent inward, with a single water molecule bridging β1 and β8, resulting in a more closed conformation consistent with the absence of a ligand or binding partner (**Figure 3H**). In contrast, the glycocholate-bound Vp-ToxSp structure adopts a more open architecture, with two waters occupying the space between the β1 and β8 strands. This arrangement parallels the Vc-ToxSp-ToxRp complex, in which Thr279 of ToxR occupies a position analogous to the two waters observed in glycocholate-bound Vp-ToxSp (**Figure 3H**). These similarities suggest that bile salt binding primes ToxS for interaction with ToxR by inducing a more open conformation of the β-barrel. This observation is also consistent with prior biochemical evidence that bile salts and detergents such as Triton X-100 enhance ToxRp-ToxSp association [18]. Together, these structural and biochemical observations indicate that bile salt binding not only drives ToxS dimerization but also prepares ToxS for a productive engagement with ToxR, thereby enabling ToxRS complex formation.

## DISCUSSION

In this study, we report the crystal structures of ToxSp in apo and glycocholate-bound forms and propose that bile-binding induces dimerization of ToxS and formation of a ToxRS heterotetramer. While ToxS is characterized as a member of the lipocalin protein family, analysis of the ToxSp topology and structure shows that it closely resembles a group of chaperone proteins including HslJ (PDB ID: 2KTS), META1, and MxiM [7,44,45,47,56]. Like ToxS, some of these proteins can stabilize their partner complexes and respond to stress. HslJ is expressed and secreted into the periplasm under stress conditions [46,52]. MxiM acts as a pilot protein for the type III secretion system in *Shigella* spp., whereas META1 is thought to play a role in secretion in *Leishmania* spp. [44,45,47]. These proteins share structure, topology, periplasmic localization of the bacterial proteins, and chaperone-like features, which suggest that ToxS and other coTTRs are evolutionarily related to a broader class of chaperone-like proteins with distinct cellular roles [12].

In addition to the topological analysis, the sequence conservation analysis of the ToxRS periplasmic domains shows that ToxS is more conserved across species than ToxR. Nonetheless, our analyses support the idea that the ToxRS system has co-evolved (**Figure S1, S2**). The most highly conserved ToxS residues are located in the hydrophobic core, proximal to the membrane, suggesting that maintaining a stable, broken β-barrel fold is crucial for ToxS function. In contrast, while the specific residues surrounding the ligand-binding pocket vary, their hydrophobic and hydrophilic natures are preserved across the Vibrionaceae family (**Table S2**). We contend that this allows the potential binding of various hydrophobic compounds, as demonstrated by: ToxR being activated by several bile salts, as well as sodium dodecyl sulfate, and the interactions between ToxRp and ToxSp increasing in the presence of Triton X-100 [18,37].

A key mechanistic question regarding the function of coTTR systems is how environmental signals activate critical transcriptional responses at the molecular level. To date, it has been proposed that the periplasmic domain of the ToxRS system senses environmental stimuli and transduces signals through the periplasmic binding interface to the cytoplasmic DNA-binding domain [7,13]. Specifically, for both ToxRS and VtrAC, it was thought that the ToxRS heterodimer potentially acted by binding to bile salts [7,11]. The structure of GCH-bound ToxSp provides the first mechanistic evidence of a physical interaction of ToxS with host-stimuli and shows ToxRS heterodimerization is not required for bile recognition. Given ligands and swapped strands are absent in the monomeric apo form, bile salt binding appears to be the driving force for ToxS dimerization.

To investigate how bile-binding could locally cluster ToxR, we modeled the ToxR periplasmic domain onto the GCH-ToxSp dimer structure. We aligned two copies of ToxRp from the *V. cholerae* heterodimeric ToxRS periplasmic domain structure onto each chain in our GCH-ToxSp structure [11]. This alignment shows that all four N-termini of the subunits appear to face the same direction and that both domain pairs are positioned to be near the membrane. Furthermore, our predicted model indicates that there are minimal steric clashes between ToxS and ToxR in the heterotetramer (**Figure 4**).

**Figure 4.**
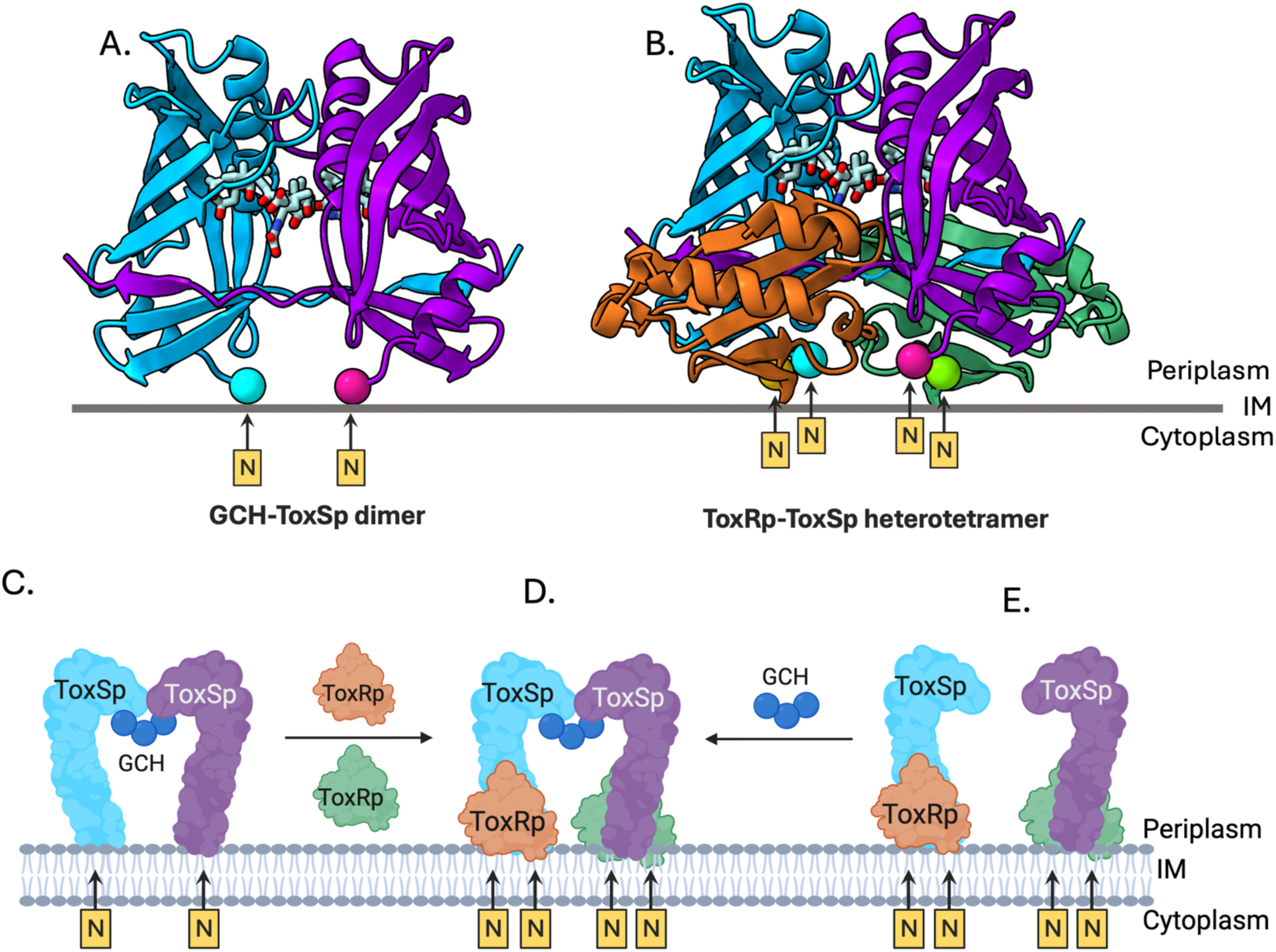
Model of the ToxRS heterotetramer in the inner membrane. (A) Three molecules of glycocholate (GCH, blue) bind to two Vp-ToxSp monomers (blue and purple β-barrels), promoting formation of a ligand-bound homodimer, anchored in the inner membrane. (B) The ToxRp-ToxSp heterotetramer model was generated using (ChimeraX) by positioning two chains of *Vibrio cholerae* ToxRp (PDB: 8ALO) onto the glycocholate-bound Vp-ToxSp dimer. The arrangement places the periplasmic domains of ToxR and ToxS in a physiologically plausible orientation with all four N-termini facing the inner membrane and minimal steric interference between ToxR and ToxS (inset). (C-E) Model of the bile-mediated ToxRS heterotetramer formation. Bile salts activate ToxRS through two possible pathways. In the first, (C) binding of three GCH molecules to monomeric apo ToxS induces formation of a domain-swapped ToxS homodimer, stabilizing an open β-barrel conformation that (D) enables ToxR recruitment and assembly of the active heterotetramer. In the second pathway, (E) bile salts engage a pre-existing apo ToxRS heterodimer, triggering conformational changes similar to those in ligand-bound ToxS that promote tetramerization of the complex. Both mechanisms converge on formation of a bile-stabilized ToxRS heterotetramer (D).

Based on the results presented here and the ToxRS model described above, we propose a mechanism for which bile salts result in the dimerization of the ToxRS complex. First, monomeric ToxS on the membrane binds bile salts, inducing dimerization as well as a conformational change in the β-barrel, opening the space between β1 and β8, making it easier for ToxR to bind ToxS (**Figure 4, path C-D**). Alternatively, if ToxS and ToxR can form a heterodimer in the absence of bile salts, we propose that bile salt binding to ToxS will subsequently induce formation of the ToxRS heterotetramer (**Figure 4, path E-D**). Overall, our model provides a scenario that suggests how bile salts may function as molecular signals that pair periplasmic sensing to activation of the ToxRS system. More generally, we propose the binding partner proteins of coTTR systems have evolved as chaperones that stabilize their cognate transcription factors and serve as binding partners mediating dimerization.

## MATERIALS AND METHODS

### Cloning of *V. parahaemolyticus* ToxS periplasmic domain

The DNA sequence of the *V. parahaemolyticus* ToxS periplasmic domain (residues S24-N171) [WP_020840240] was codon-optimized and synthesized to include a SapI site, the coding sequence, and a BamHI site, flanked by primer binding sites. The fragment was amplified with PCR, digested with SapI and BamHI restriction enzymes, and purified using the PCR purification kit (Qiagen). The pTYB21 vector (NEB), containing an N-terminal chitin-binding domain intein (CBD-intein) affinity tag, was digested with the same restriction enzymes (SapI and BamHI), treated with calf intestinal phosphatase (CIP; NEB), and purified using the PCR purification kit (Qiagen) (**Table S6**). The ToxSp insert was ligated into the prepared pTYB21 vector using Quick Ligase (NEB), and the ligation products were transformed into *E. coli* DH5α cells (NEB). The transformants were selected on an LB agar plate containing 100 µg/mL carbenicillin and screened by colony PCR for the correct insert size. The selected colonies were grown in small-scale cultures; a portion was stored at -80 °C, while plasmid DNA from the remaining culture was purified using the Miniprep kit (Qiagen). Plasmid sequences were confirmed by Sanger sequencing with primers complementary to the T7 promoter.

### Protein Expression and Purification

The confirmed CBD-ToxSp plasmid was transformed into *E. coli* BL21-CodonPlus (DE3) competent cells (Agilent Technologies). The starter culture was grown overnight at 30 °C in ZYP-0.8G medium supplemented with 200 µg/mL carbenicillin and 25 µg/mL chloramphenicol in a shaking incubator [57]. The culture was used to inoculate Terrific Broth Modified media (Fisher Scientific) at a 1:250 ratio, supplemented with 2 mM MgSO_4_, 200 µg/mL carbenicillin, and 25 µg/mL chloramphenicol. The cells were grown at 37 °C to an OD_600_ of 2-3, induced with 1 mM IPTG, and incubated at 18 °C for 16-20 hours. The cells were harvested by centrifugation at 4,500 × *g* for 25 minutes at 4 °C and resuspended in lysis buffer (20 mM Tris-HCl pH 8.0, 200 mM NaCl, 1 mM EDTA) containing cOmplete protease inhibitor cocktail tablets (Roche). Lysis was performed by passing the cells through a French press three times, and the lysate was centrifuged at 100,000 × *g* for 45 minutes at 4 °C. The supernatant was filtered through a 0.45 µm filter before purification. The supernatant was passed over chitin-binding resin (NEB) pre-equilibrated with Tris wash buffer (20 mM Tris-HCl pH 8.0, 200 mM NaCl) to capture the CBD-ToxSp fusion protein. The resin was washed sequentially with 10 column volumes (CV) of high-salt Tris wash buffer (20 mM Tris-HCl pH 8.0, 1 M NaCl) and 20 CV of Tris wash buffer. The resin was rapidly washed with 3 CV of Tris cleavage buffer (20 mM Tris-HCl pH 8.0, 200 mM NaCl, 50 mM DTT, 0.5 mM PMSF, 1 mM EDTA, cOmplete protease cocktail tablet) to saturate the resin and initiate on-column intein-mediated cleavage of the CBD tag. The column was incubated at 23 °C for 16-40 hours. Cleaved ToxSp protein was eluted in Tris wash buffer, concentrated using Amicon Ultra 3 kDa molecular weight cut-off (MWCO) centrifugal filters (Millipore), and further purified by size-exclusion column (HiLoad 16/600 Superdex 200 pg, Cytiva) pre-equilibrated with gel filtration buffer (20 mM Tris-HCl pH 8.0, 200 mM NaCl). The purified ToxSp protein solution was concentrated to 9-16 mg/mL, flash-frozen in liquid nitrogen, and stored at -80 °C until crystallization.

### Crystallization and Data Collection

To obtain apo ToxSp crystals, a seeded crystallization method was used to promote the growth of diffraction-quality crystals. First, purified ToxSp protein was concentrated to 16 mg/mL and used to generate initial crystals. These initial ToxSp crystals were grown by combining protein and crystallization solution (100 mM succinic acid pH 6.0, 16% (w/v) PEG 3,350) at a 1:1 ratio in a 96-well, 2-drop MRC plate (Swissci) with NT8 Drop Setter (Formulatrix). The resulting crystals were harvested and stored in 50 µL crystallization solution, then disrupted with a polytetrafluoroethylene seed bead (Hampton Research) to generate a seed stock. This stock solution was serially diluted (1:10 to 1:10,000) and stored at room temperature (23 °C). For seeded crystallization, purified ToxSp protein was concentrated to 14 mg/mL and seeded by adding the diluted seed stock at a 25:2 (v/v) protein-to-seed ratio. The crystallization trials were set up in the 96-well, 2-drop MRC plates (Swissci) using the NT8 Drop Setter (Formulatrix) with protein-to-reservoir ratios of 1:1 (drop 1) and 1:2 (drop 2), using 96-well sparse matrix crystal screens. Under these conditions, the optimal diffraction-quality apo crystals were obtained from Berkeley crystal screen condition E8 (Rigaku Reagents; 100 mM HEPES pH 7.5, 200 mM sodium malonate dibasic pH 7.0, 20% (w/v) PEG monomethyl ether 2,000). The crystals were cryoprotected with a solution formulated with 70% crystal reservoir solution and 30% glycerol. The diffraction data were collected at the NSLS-II AMX beamline (Brookhaven National Laboratory) and processed to 2.1 Å resolution with XDS, yielding a space group of P 2_1_ 2_1_ 2_1_ with unit cell parameters 53.746 58.230 89.456 90.000 90.000 90.000 (**Table S1**) [58].

To obtain glycocholate-bound ToxSp crystals, the frozen ToxSp protein stock at 16 mg/mL was thawed and supplemented with 10 mM sodium glycocholate stock prepared in ToxSp buffer (20 mM Tris-HCl pH 8.0, 200 mM NaCl) to achieve final concentrations of approximately 622 µM ToxSp and 3.13 mM sodium glycocholate (1:5 protein-to-ligand ratio). The combined glycocholate-ToxSp solution was incubated overnight at 4 °C with rotation to allow complex formation. The crystallization plates for the glycocholate-bound complex were set up using the same seeding method as for apo ToxSp. The glycocholate-ToxSp solution was seeded by adding diluted seed stock prepared from apo ToxSp crystals at a 25:2 (v/v) protein-to-seed ratio. The crystallization trials were set up in 96-well, 2-drop MRC plates (Swissci) using 96-well sparse matrix crystal screens. The initial crystal hit was observed from Index crystal screen condition H9 (Hampton Research). The Index H9 condition was optimized to 0.06 M zinc acetate dihydrate and 17% (w/v) PEG 3,350. The crystals were cryoprotected with a solution formulated with 70% crystal reservoir solution and 30% glycerol. The diffraction data were processed to 1.9 Å resolution with autoPROC, yielding a space group of P 2_1_ 2_1_ 2_1_ and a unit cell of 69.828 92.503 95.805 90.00 90.00 90.00 (**Table S1**) [59].

### Structure Determination

The structure of apo ToxSp was determined by molecular replacement using an AlphaFold structure prediction model, truncated to exclude regions with pLDDT under 90 [60]. The molecular replacement was performed using Phaser [61]. The iterative refinement cycles were performed in Refine from PHENIX, and manual model building was performed in COOT [62]. Structural visualization and analysis were performed in ChimeraX [63]. The final structure was deposited in the Protein Data Bank (PDB ID: 8U2F). The strand-swapped dimer structure of GCH-bound ToxSp was solved by molecular replacement using a RoseTTAFold All-Atom prediction model generated as a monomer [64]. The electron density map clearly indicated a strand swap in the β8 region, confirmed by a composite omit map in PHENIX [65]. The protein structure and GCH ligands were built using COOT with ligand restraints from the REFMAC library [62,66]. The final structure visualization and analysis were performed in ChimeraX [63].

### Conservation analysis

To analyze the ToxSp sequence conservation, BLAST was used to find sequences homologous to the *V. cholerae* ToxR periplasmic domain residues 199-294 [67,68]. The resulting BLAST sequences were then curated by removing entries that were multispecies alignments, partial sequences, identical sequences, or sequences from the same species that were highly similar, leaving a list of ToxR periplasmic domains primarily representing a single sequence per species. The curated ToxRp sequences were aligned using MUSCLE and then used to generate a percent identity matrix with Clustal Omega [41,69,70]. Based on this ToxRp sequence alignment, the corresponding ToxSp sequences from the same species were identified in the NCBI and compiled into the same order as the aligned ToxRp sequences to preserve one-to-one pairing [68]. This procedure resulted in a list of 280 paired ToxR and ToxS sequences. The ToxS sequences were then aligned, and the sequences of the N-terminal tail and transmembrane domain were removed so that only periplasmic domains (ToxSp) were retained. Both the ToxRp and ToxSp sequences were subsequently aligned again, and a percent identity matrix calculated by Clustal Omega without rearrangement of the sequence order [69]. Once the alignment was complete, the file with the ToxSp alignment was loaded into ChimeraX to calculate and visualize the protein conservation at the residue level using the built-in program AL2CO [42].

## ADDITIONAL INFORMATION

Atomic coordinates and structure factors for the reported structures have been deposited in the Protein Data Bank (PDB) under accession numbers 8U2F (apo Vp-ToxSp) and 9ZJO (GCH-bound Vp-ToxSp). The data collection statistics are shown in Table S1 in supplementary information.

## ACKNOWLEDGEMENTS

This work was supported by National Institute of Allergy and Infectious Diseases (NIAID) R01AI168157 and R21AI140740 to JFK. In addition, this work was also supported by a National Science Foundation (NSF) CAREER award (#2045671), a Burroughs Wellcome Investigator in the Pathogenesis of Infectious Disease (#1021977) and ALSAC at St. Jude Children’s Research Hospital to SAM.

