## Supplemental tables and figures for "Bile-mediated ToxS homodimerization provides a model for ToxRS periplasmic interactions"

Running title: Bile mediates ToxS periplasmic domain homodimerization

**Supplemental Information**

**Table S1. Data collection and refinement statistics for ToxSp.**

| | **Vp-ToxSp** (PDB ID: 8U2F) | **GCH bound Vp-ToxSp** (PDB ID: 9ZJO) |
|---|---|---|
| **Wavelength (Å)** | 0.920112 | 0.920105 |
| **Resolution range (Å)** | 26.92 - 2.004 (2.076 - 2.004) | 66.55 - 1.961 (2.01 - 1.96) |
| **Space group** | P 21 21 21 | P 21 21 21 |
| **Unit cell (Å, º)** | 53.85 58.33 89.68 90 90 90 | 69.828 92.503 95.805 90 90 90 |
| **Total reflections** | 172599 (17411) | 274081 (18185) |
| **Unique reflections** | 19566 (1912) | 44411 (2966) |
| **Multiplicity** | 8.8 (9.1) | 6.2 (6.1) |
| **Completeness (%)** | 99.84 (99.69) | 97.63 (98.15) |
| **Mean I/sigma(I)** | 10.99 (0.94) | 6.71 (1.06) |
| **Wilson B-factor (Å²)** | 43.25 | 27.88 |
| **R-merge** | 0.1052 (1.984) | 0.207 (2.569) |
| **R-meas** | 0.112 (2.103) | 0.2264 (2.809) |
| **R-pim** | 0.03781 (0.6912) | 0.09038 (1.124) |
| **CC1/2** | 0.998 (0.526) | 0.995 (0.439) |
| **CC*** | 1 (0.83) | 0.999 (0.781) |
| **Reflections used in refinement** | 19551 (1912) | 44103 (2911) |
| **Reflections used for R-free** | 1958 (191) | 2160 (147) |
| **R-work** | 0.2252 (0.3039) | 0.2013 (0.2717) |
| **R-free** | 0.2711 (0.3369) | 0.2343 (0.3098) |
| **CC(work)** | 0.938 (0.769) | 0.959 (0.759) |
| **CC(free)** | 0.897 (0.748) | 0.974 (0.741) |
| **Number of non-hydrogen atoms** | 2334 | 5591 |
| **macromolecules** | 2243 | 4772 |
| **ligands** | 0 | 271 |
| **solvent** | 91 | 548 |
| **Protein residues** | 281 | 596 |
| **RMS(bonds) (Å)** | 0.010 | 0.009 |
| **RMS(angles) (°)** | 1.38 | 1.07 |
| **Ramachandran favored (%)** | 100.00 | 98.13 |
| **Ramachandran allowed (%)** | 0.00 | 1.87 |
| **Ramachandran outliers (%)** | 0.00 | 0 |
| **Rotamer outliers (%)** | 0.38 | 0.71 |
| **Clashscore** | 13.77 | 5.31 |
| **Average B-factor (Å²)** | 50.02 | 33.91 |
| **macromolecules** | 49.89 | 31.92 |
| **ligands** | | 58.99 |
| **solvent** | 53.09 | 38.86 |

**Table S2. Sequence alignment and chemical properties of ToxSp binding pocket from 280 Vibrionaceae species.**

| ToxSp Seq. Alignment Position | Vp-ToxSp Seq. Position | Vp-ToxSp AA | % Identity | % Hydrophobic | % Hydrophilic | % Charged | % Aromatic | % Pro/Gly |
|---|---|---|---|---|---|---|---|---|
| 43 | 41 | Met (M) | 53.93 | 66.43 | 32.86 | 0.71 | 0 | 0 |
| 45 | 43 | Thr (T) | 57.86 | 13.21 | 82.50 | 0.71 | 1.43 | 1.79 |
| 47 | 45 | Ile (I) | 74.64 | 97.86 | 0.36 | 0.36 | 1.43 | 0 |
| 49 | 47 | Asp (D) | 26.43 | 9.29 | 25.36 | 51.07 | 2.86 | 8.93 |
| 51 | 49 | Leu (L) | 21.79 | 36.79 | 21.07 | 33.93 | 0.71 | 5.00 |
| 57 | 50 | Pro (P) | 7.86 | 17.14 | 43.57 | 13.93 | 9.64 | 8.93 |
| 64 | 54 | Val (V) | 33.93 | 70.00 | 6.79 | 1.43 | 0.71 | 0 |
| 70 | 57 | Leu (L) | 92.86 | 99.64 | 0.36 | 0 | 0 | 0 |
| 73 | 60 | Val (V) | 68.93 | 93.57 | 5.71 | 0 | 0 | 0.71 |
| 75 | 62 | Val (V) | 37.86 | 75.71 | 19.29 | 3.57 | 0.71 | 0.71 |
| 91 | 76 | Arg (R) | 90.36 | 0.71 | 0 | 99.29 | 0 | 0 |
| 95 | 80 | Ile (I) | 22.5 | 99.29 | 0 | 0 | 0.71 | 0 |
| 97 | 82 | Leu (L) | 86.79 | 95.00 | 0 | 0 | 4.64 | 0 |
| 116 | 93 | Ile (I) | 71.07 | 98.57 | 0 | 0 | 1.43 | 0 |
| 158 | 134 | Ile (I) | 80 | 100.00 | 0 | 0 | 0 | 0 |
| 161 | 137 | Ile (I) | 28.93 | 77.50 | 0 | 1.43 | 20.71 | 0 |
| 162 | 138 | Phe (F) | 67.5 | 22.50 | 0 | 0 | 77.50 | 0 |
| 165 | 141 | Asp (D) | 73.57 | 0.71 | 13.93 | 80.36 | 0 | 4.64 |

To generate the percent identity, the frequency of each amino acid in the alignment was determined using UGENE [1]. Applying a STATA (StataCorp) script, the frequency was used to calculate the percentage of each amino acid at each position. To obtain the percentages of hydrophobic, hydrophilic, charged, aromatic, and pro/gly the percentages of the selected amino acids for each category added together.

1. Okonechnikov K, Golosova O, Fursov M, Varlamov A, Vaskin Y, Efremov I, et al. Unipro UGENE: a unified bioinformatics toolkit. Bioinformatics. 2012;28: 1166–1167. doi:10.1093/BIOINFORMATICS/BTS091

**Table S3. DALI search result showing Vp-ToxSp structural similarity with chaperone proteins.**

| No | PDB ID-Chain | Z | rmsd | lali | nres | %id PDB | Description |
|---|---|---|---|---|---|---|---|
| 1 | 5kew-B | 10.6 | 2.6 | 109 | 132 | 9 PDB | MOLECULE VTRA PROTEIN; |
| 2 | 4nyq-A | 9.3 | 2.7 | 98 | 153 | 9 PDB | MOLECULE MILK PROTEIN; |
| 3 | 4u3q-B | 9.2 | 2.1 | 86 | 99 | 17 PDB | MOLECULE 17 KDA LIPOPROTEIN; |
| 4 | 2kts-A | 8.8 | 3.1 | 97 | 117 | 7 PDB | MOLECULE HEAT SHOCK PROTEIN HSLJ; |
| 5 | 2la7-A | 8.1 | 2.8 | 103 | 145 | 13 PDB | MOLECULE UNCHARACTERIZED PROTEIN; |
| 6 | 3lhn-A | 8 | 2.7 | 91 | 108 | 8 PDB | MOLECULE LIPOPROTEIN; |
| 7 | 4n7c-A | 7.7 | 2.8 | 100 | 174 | 9 PDB | MOLECULE BLA G 4 ALLERGEN VARIANT 1; |
| 8 | 2erv-A | 7.7 | 3.1 | 98 | 150 | 3 PDB | MOLECULE HYPOTHETICAL PROTEIN PAER03002360; |
| 9 | 4kh8-A | 7.7 | 2.4 | 90 | 311 | 10 PDB | MOLECULE HYPOTHETICAL PROTEIN; |
| 10 | 4rlc-A | 7.7 | 3.2 | 102 | 135 | 7 PDB | MOLECULE OUTER MEMBRANE PORIN F; |
| 11 | 4iab-A | 7.5 | 4.2 | 95 | 142 | 13 PDB | MOLECULE HYPOTHETICAL PROTEIN; |
| 12 | 2jw1-A | 7.5 | 3.1 | 93 | 115 | 13 PDB | MOLECULE LIPOPROTEIN MXIM; |
| 13 | 2mhd-A | 7.5 | 3 | 90 | 110 | 11 PDB | MOLECULE UNCHARACTERIZED PROTEIN; |
| 14 | 8ehd-A | 7.4 | 3.4 | 82 | 101 | 13 PDB | MOLECULE POTEMPIN E (POTE); |
| 15 | 7b2a-A | 7.3 | 2.5 | 92 | 162 | 7 PDB | MOLECULE CIRPA5; |
| 16 | 1qft-A | 7.3 | 2.7 | 95 | 175 | 7 PDB | MOLECULE PROTEIN (FEMALE-SPECIFIC HISTAMINE BINDING |
| 17 | 4lqz-A | 7.2 | 2.6 | 83 | 131 | 8 PDB | MOLECULE UNCHARACTERIZED PROTEIN; |
| 18 | 2z4h-B | 7.1 | 2.8 | 82 | 194 | 13 PDB | MOLECULE COPPER HOMEOSTASIS PROTEIN CUTF; |
| 19 | 5hcc-C | 7 | 2.5 | 90 | 148 | 6 PDB | MOLECULE COMPLEMENT C5; |
| 20 | 2m4l-A | 6.9 | 3.4 | 82 | 99 | 6 PDB | MOLECULE PROTEIN BT_0846; |
| 21 | 1i4u-A | 6.9 | 2.6 | 90 | 181 | 10 PDB | MOLECULE CRUSTACYANIN; |
| 22 | 3rby-A | 6.9 | 3.4 | 95 | 245 | 8 PDB | MOLECULE UNCHARACTERIZED PROTEIN YLR301W; |
| 23 | 4tq2-A | 6.9 | 2.8 | 91 | 177 | 12 PDB | MOLECULE PUTATIVE PHYCOERYTHRIN LYASE; |
| 24 | 6x1k-A | 6.9 | 3.4 | 97 | 124 | 7 PDB | MOLECULE DE NOVO DESIGNED TRANSMEMBRANE BETA-BARREL TMB2.3 |
| 25 | 1jmx-A | 6.8 | 3.5 | 96 | 493 | 7 PDB | MOLECULE AMINE DEHYDROGENASE; |

**Table S6. Sequence and primers for ToxS constructs.**

| Protein | Vector | Amino acid sequence |
|---|---|---|
| Vp-ToxSp | pTYB21 | MSDLKVEQVLTSNEWQSTMVTVITDNLPDDTVGPLRRVNVESNVKYLPNGDYIRVSNIKLFAQGSTAE STINISEKGRWEVSDNYLLVSPSEFKDISSSQSKDFSEAQLRLITQIFKLDAEQSRRIDVVNEKTLLLTSL NHGSTVLFRN |

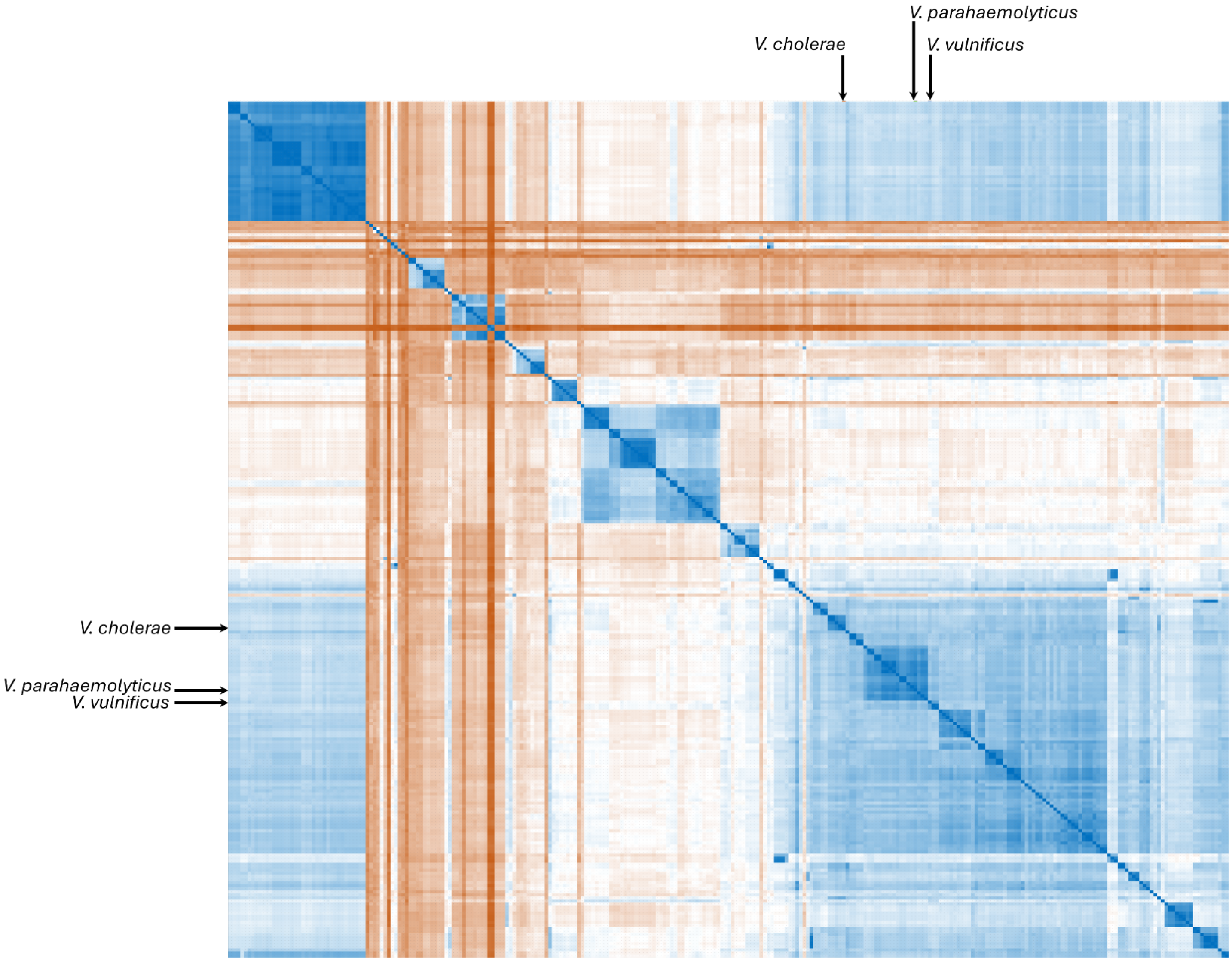


**Figure S1. Pairwise percent identity matrix for ToxSp sequence across 280 sequences from Vibrionaceae family.** Each pixel shows the pairwise sequence percent identity generated from multiple sequence alignment between 280 Vibrionaceae ToxSp sequences. The diagonal (dark blue) corresponds to self-comparison (100% identity). The rows and columns represent same order of sequences matched pairwise to create a symmetric matrix. The order of the 280 sequences is the same as that of ToxRp matrix for direct comparison. The color indicates percent identity (blue, high identity; white, intermediate identity; orange, low identity). A higher proportion of blue and white pixels in ToxSp matrix compared to the ToxRp matrix shows that ToxSp sequences are generally more similar to one another than ToxRp.

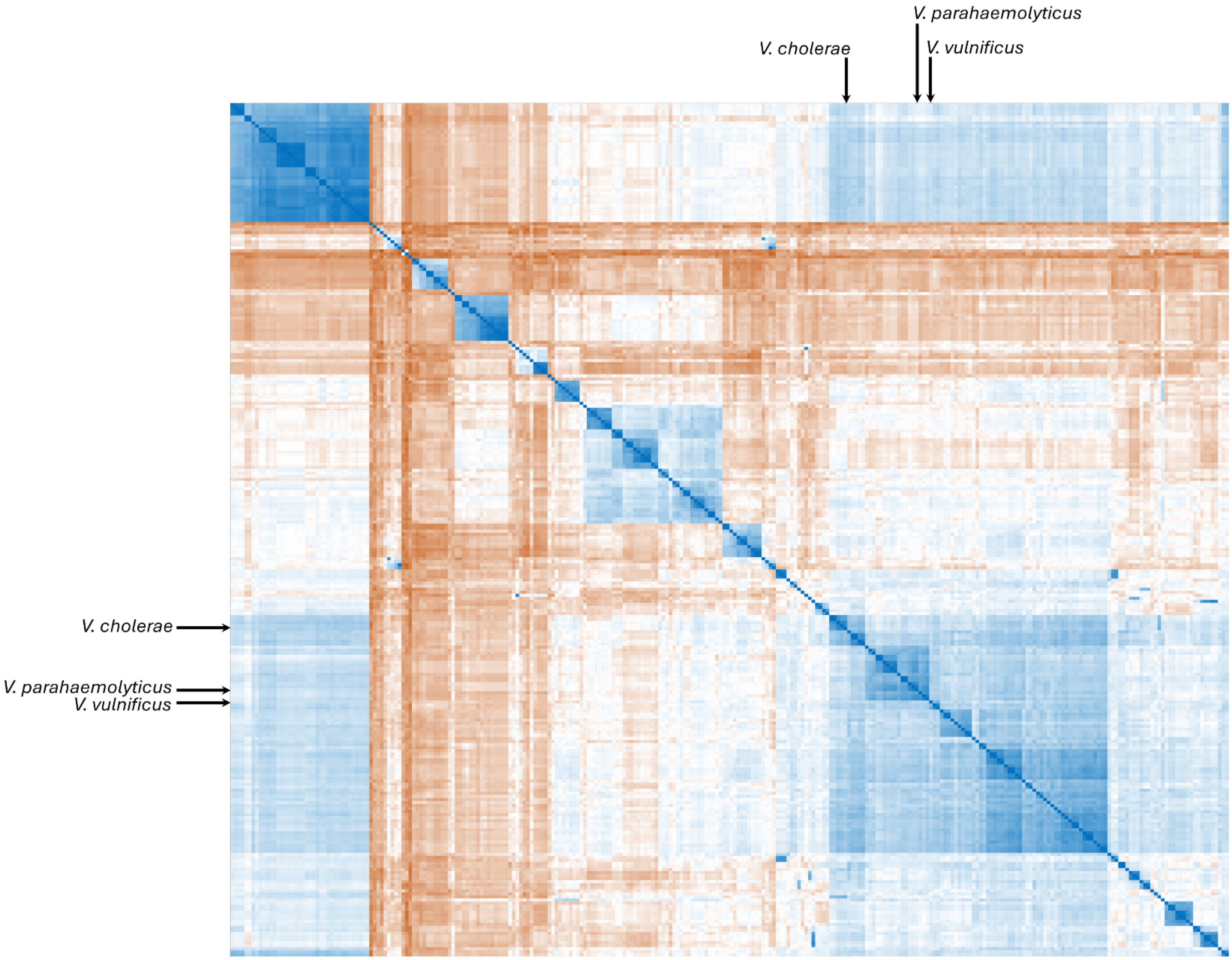


**Figure S2. Pairwise percent identity matrix for ToxRp sequence across 280 sequences from Vibrionaceae family in the same order as ToxSp matrix.** Each pixel shows the pairwise sequence percent identity generated from multiple sequence alignment between 280 Vibrionaceae ToxRp sequences. The diagonal (dark blue) corresponds to self-comparison (100% identity). The rows and columns represent same ordered list of sequences matched pairwise to create a symmetric matrix. The order of the 280 sequences is the same as that of ToxSp matrix for direct comparison. The color indicates percent identity (blue, high identity; white, intermediate identity; orange, low identity). A higher proportion of orange pixels and fewer blue pixels in the ToxRp matrix compared to ToxSp indicates that ToxRp is less conserved than ToxSp.

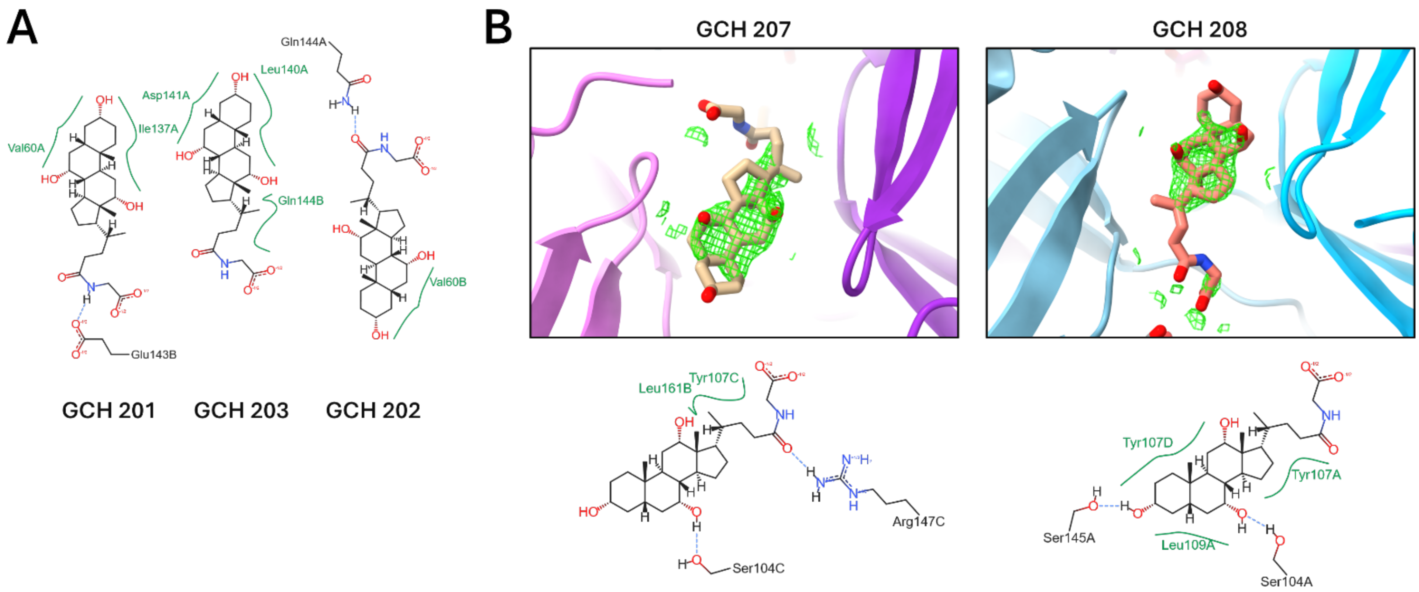


**Figure S3. Glycocholate molecules observed at the binding pocket and at the crystal packing sites.** (**A**) 2D interaction diagram of the three glycocholate molecules bound between the two chains of the strand-swapped dimer (GCH 201-203) with nearby interacting residues shown. (**B**) Two additional glycocholate molecules observed at the crystal-packing interfaces (GCH 207-208), which are the peripherally bound ligands between the two dimer chains in the crystallographic lattice. 2D interaction diagram of GCH 207-208 are shown alongside the composite omit electron density (green mesh; contoured at 1 σ). Density is the strongest at the ring group and weaker at the glycine-conjugated tail. These ligands are not interpreted as physiologically relevant binding events.
